# Induction of cross-reactive antibody responses against the RBD domain of the spike protein of SARS-CoV-2 by commensal microbiota

**DOI:** 10.1101/2021.08.08.455272

**Authors:** Justus Ninnemann, Lisa Budzinski, Marina Bondareva, Mario Witkowski, Stefan Angermair, Jakob Kreye, Pawel Durek, S. Momsen Reincke, Elisa Sánchez-Sendin, Selin Yilmaz, Toni Sempert, Gitta Anne Heinz, Caroline Tizian, Martin Raftery, Günther Schönrich, Daria Matyushkina, Ivan V. Smirnov, Vadim M. Govorun, Eva Schrezenmeier, Thomas Dörner, Silvia Zocche, Edoardo Viviano, Katharina Johanna Sehmsdorf, Hyun-Dong Chang, Philipp Enghard, Sascha Treskatsch, Andreas Radbruch, Andreas Diefenbach, Harald Prüss, Mir-Farzin Mashreghi, Andrey A. Kruglov

## Abstract

The commensal microflora is a source for multiple antigens that may induce cross-reactive antibodies against host proteins and pathogens. However, whether commensal bacteria can induce cross-reactive antibodies against SARS-CoV-2 remains unknown. Here we report that several commensal bacteria contribute to the generation of cross-reactive IgA antibodies against the receptor-binding domain (RBD) of the SARS-CoV-2 Spike protein. We identified SARS-CoV-2 unexposed individuals with RBD-binding IgA antibodies at their mucosal surfaces. Conversely, neutralising monoclonal anti-RBD antibodies recognised distinct commensal bacterial species. Some of these bacteria, such as *Streptococcus salivarius*, induced a cross-reactive anti-RBD antibodies upon supplementation in mice. Conversely, severely ill COVID-19 patients showed reduction of *Streptococcus* and *Veillonella* in their oropharynx and feces and a reduction of anti-RBD IgA at mucosal surfaces. Altogether, distinct microbial species of the human microbiota can induce secretory IgA antibodies cross-reactive for the RBD of SARS-CoV-2.

## Main text

SARS-CoV-2 virus infects cells via interaction of the Spike (S) protein with the ACE2 receptor, which is expressed by various cell types [1, 2, 3]. The Spike protein of SARS-CoV-2 contains a receptor-binding domain (RBD) that mediates its interaction with ACE2 and viral entry [3, 4]. Blocking of this crucial interaction by monoclonal anti-SARS-CoV-2-RBD antibodies confers protection of the host against infection of target cells [5, 6]. Systemically distributed antibodies (mainly IgG, IgM, and IgA1) curtail virus propagation after productive infection of the host, while the presence of antigen-specific antibodies secreted at the mucosal surfaces (IgA2, IgA1, and IgM) may prevent initial infection of the host [7]. The absence of IgA2 antibodies specific for SARS-CoV-2 antigens in severely diseased COVID-19 patients has also been demonstrated [8], suggesting that mucosal anti-viral IgA antibodies may protect the host from a severe course of COVID-19. Several studies have reported the presence of RBD-binding antibodies in unexposed healthy individuals [9, 10, 11, 12, 13]. Induction of such antibodies by previous infections with common cold coronaviruses has been postulated, but this link has not been formally proven. The original antigens inducing cross-reactive RBD-binding secretory IgA antibodies have remained obscure.

IgA antibodies at mucosal surfaces are mainly induced by commensal microbiota [14]. It is estimated that the human microbiota contains several millions of genes [15], thus potentially providing a plethora of epitopes for antibodies [16]. Some of such epitopes may resemble host proteins, potentially inducing autoimmunity [17, 18, 19, 20, 21], while others may resemble proteins from other microorganisms and mediate cross-reactive immunity [17, 22]. Microbiota-induced cross-reactive immunity also provides protection against microbial infections by *Citrobacter rodentium, Clostridiodes difficile, Pseudomonas aeruginosa* [23] and by viruses like influenza [24]. Protection is mediated by increasing fitness of the innate immune system, e.g. via tonic type I IFN production [25, 26], and by cross-reactive adaptive antibody responses [23]. Interestingly, cross-reactive antibodies targeting gp41 of HIV-1 are induced by commensal microbiota [27]. Here we describe the induction of cross-reactive antibody responses targeting SARS-CoV-2 by distinct members of the oral and gut microbiota.

We initially had analysed RBD-specific IgA in the fecal supernatants of age-matched healthy individuals and severely diseased COVID-19 patients (Table S1). Two out of 12 age-matched healthy donors, previously unexposed to SARS-CoV-2, as confirmed by lack of anti-NP SARS-CoV-2 IgG antibodies in their sera (Fig. S1A), did have fecal IgA antibodies reactive to RBD (Fig. 1A), 10 out of 21 severely diseased COVID-19 patients had fecal IgA specific for Spike protein RBD of SARS-CoV-2 (Fig. 1B and Fig. S1B). Considering that age is an important risk factor for the development of severe COVID-19, we next determined the prevalence of RBD-binding IgA antibodies in young unexposed individuals (Fig. 1C, D and Table S1). We detected RBD-binding fecal IgA in approximately 50% of young healthy donors and the magnitude of the RBD-binding IgA responses in feces negatively correlated with the age of the donors (Fig. 1E). Given the compositional complexity of fecal supernatant, we next purified IgA antibodies and tested whether the mucosal RBD-binding IgA can inhibit binding of RBD protein to the ACE2 receptor, thereby potentially blocking the entry of SARS-CoV-2 into the host cells. To this end, we expressed human ACE2 on 293T cells, then incubated the ACE2-expressing cells with biotinylated RBD in the presence of purified mucosal IgA of various healthy donors (Fig. 1F, S1C). The fraction of bound RBD was analysed by flow cytometry using fluorescent streptavidin. Purified intestinal IgA from 5 out of 14 healthy donors inhibited RBD binding to ACE2 (Fig. 1F). Of note, complete inhibition of ACE2-RBD interaction was not achieved even at 1:1 dilution, indicating a rather low concentration of neutralising anti-RBD IgA in the feces. Also, IgA from some donors with anti-RBD antibodies did not inhibit the RBD-ACE2 interaction, indicating that healthy individuals may harbor both inhibitory and non-inhibitory IgA antibodies directed against the RBD of SARS-CoV-2 (Fig. 1F). Interestingly, healthy donors exhibited IgA2 antibodies specific for RBD in their feces, while severely diseased COVID-19 patients lacked fecal anti-RBD IgA2, consistent with a previous report [8] (Fig. 1G).

**Fig. 1.**
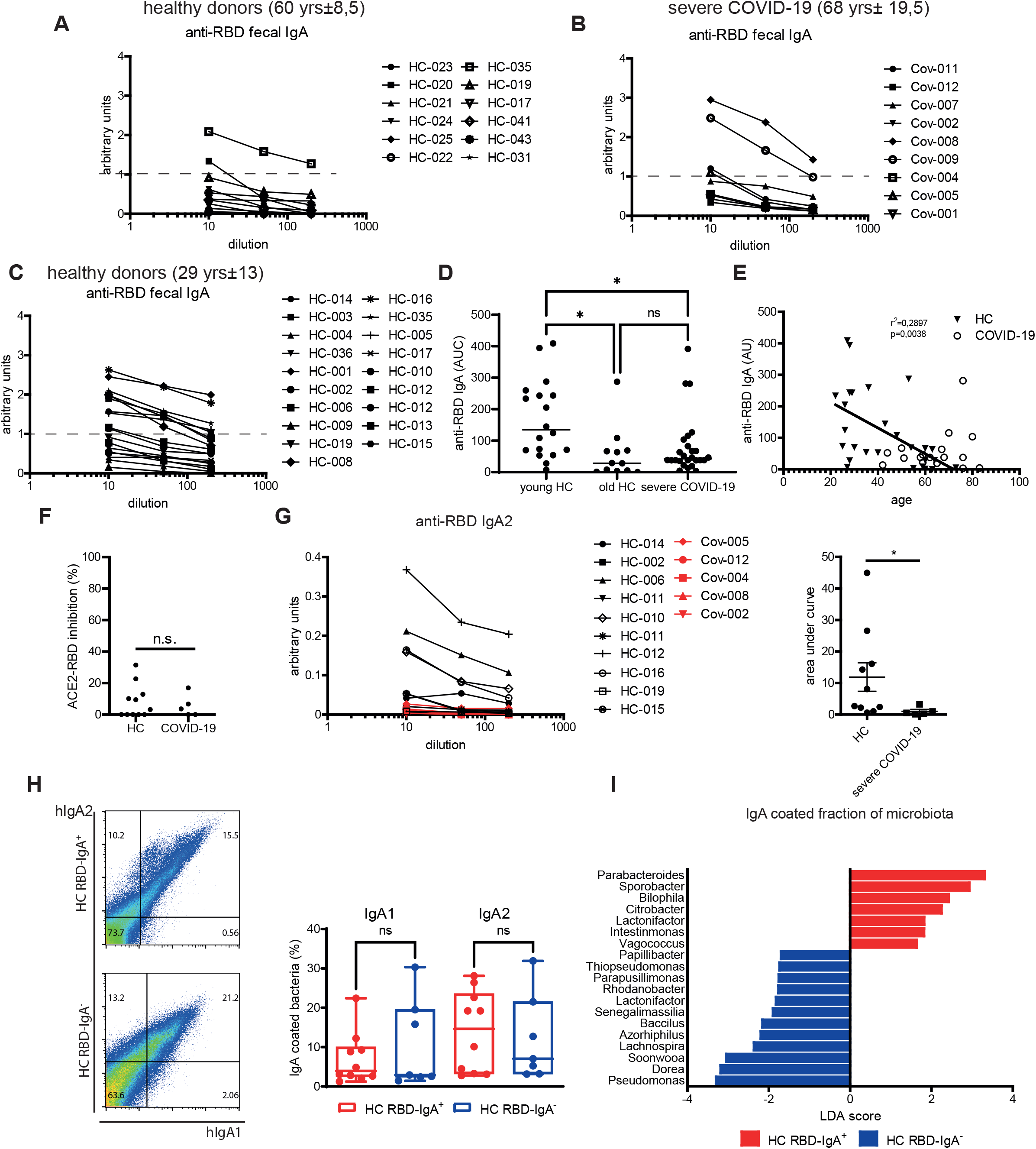
Presence of mucosal IgA antibodies reactive against RBD in healthy individuals. Levels of anti-RBD IgA in fecal supernatants of age-matched healthy (**A**) and severe COVID-19 (**B**) individuals. (**C**) Levels of anti-RBD IgA in fecal supernatants of young healthy individuals. (**D**) Area under the curve (AUC) values for the anti-RBD IgA ELISA measurement of the donors presented in (**A-C**). Correlation of the levels of anti-RBD IgA with the age in healthy individuals and COVID-19 patients. Inhibition of RBD binding to ACE2 by IgA purified from feces of healthy people and severe COVID-19 patients. (**G**). Levels and AUC values of anti-RBD IgA2 in purified IgA fraction from healthy and severe COVID-19 individuals. (**H**). Representative dot plots and quantification of fecal IgA coating from healthy individuals that have anti-RBD IgA (HC RBD-IgA+) or lack anti-RBD IgA (HC RBD-IgA-). (**I**) Linear discriminant analysis (LDA) scores of the IgA bound bacterial fraction isolated from HC RBD-IgA+ and HC RBD-IgA-. *, p<0.05, **, p<0.01, ***, p<0.001, as calculated by unpaired t-test (F, G, H) or by Kruskal-Wallis test with Dunn’s multiple comparisons (D); ns, not significant.

IgA is induced by microbiota and does bind to microbiota [28]. Thus we next analysed whether RBD-binding IgA also recognizes commensal microbiota. To this end, we first divided our healthy cohort (HC) in two groups based on the presence or absence of RBD-binding IgA in their fecal supernatants: HC RBD-IgA^+^ and HC RBD-IgA^-^, respectively, and quantified the coating of bacteria by endogenous IgA. Both donor groups exhibited similar coating of their intestinal microbiota by mucosal IgA1 and IgA2 (Fig. 1H). To identify the bacteria binding to mucosal IgA1 and IgA2, we isolated them by fluorescence-activated cell sorting and determined their taxonomic composition by 16S rRNA sequencing. Linear discriminant (LDA) combined with effect size (LefSE) analysis revealed distinct taxonomic differences of IgA coated bacteria of RBD-IgA^+^ versus RBD-IgA^-^ healthy donors. The IgA coated bacterial fraction of RBD IgA^+^ donors were enriched for *Parabacteroides, Sporobacter, Bilophila*, and *Vagococc*us, while in RBD-IgA^-^ donors the IgA coated fraction was enriched for *Pseudomonas, Dorea, Soonwooa, Lachnospira*, and *Bacillus* genera (Fig. 1I). These data suggest that mucosal anti-RBD IgA is associated with recognition of distinct commensal microbiota by mucosal IgA.

To directly test whether anti-RBD antibodies bind to commensal bacteria, we stained the fecal microbiota of healthy individuals with neutralising anti-RBD antibodies that had either been generated in immunized rabbits or that had been cloned from hospitalised COVID-19 patients [29]. The neutralising rabbit antibody showed binding to a significant fraction of microbiota from HC (Fig. 2A). Furthermore, out of 15 monoclonal neutralising antibodies derived from hospitalized COVID-19 patients (for the details see [29]) only two (HK CV07-287, HL CV07-250) showed no microbiota binding activity (Fig. 2B, C). The remaining antibodies recognised commensal bacteria, 9 of them also independently of pre-existing fecal anti-RBD IgA (Fig. 2B, C). Of note, two clonally related antibodies, CV07-200 and CV07-283, showed distinct binding patterns (Fig. 2C and Fig. S2). Co-staining of microbiota with rabbit and human monoclonal antibodies showed that both recognize similar as well as distinct fecal bacteria communities (Fig. S3). Thus, most neutralising human anti-RBD SARS-CoV-2 antibodies tested in our study bind to distinct commensal bacteria.

**Fig. 2.**
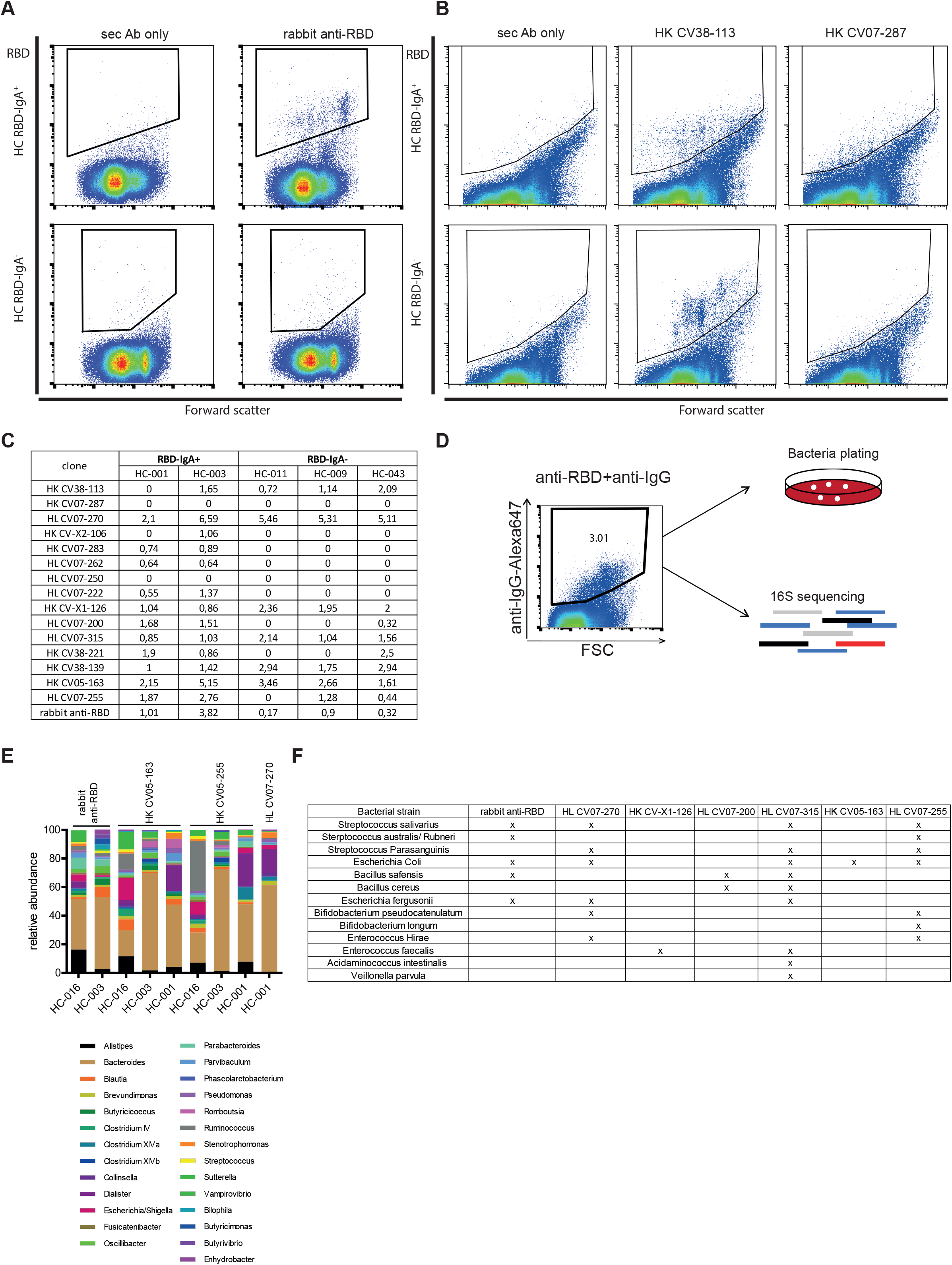
Neutralising anti-RBD antibodies recognize distinct commensal bacteria. Representative dot plots of human fecal microbiota stained with neutralising anti-RBD antibody raised in rabbit. (**B**) Representative dot plots of microbiota stained with monoclonal neutralising anti-RBD antibodies derived from COVID-19 patients. (**C**) Frequency of bacteria bound by human neutralising anti-RBD antibodies towards microbiota from healthy individuals. Fecal microbiota from 5 heathy donors were stained with 15 monoclonal anti-RBD antibodies from COVID-19 patients or anti-rabbit RBD, followed by respective secondary fluorochrome-coupled antibodies. Bound bacterial fraction was defined via comparison of stained sample with sample stained only with secondary antibody. (**D**) Strategy for the identification of bacteria that is bound by anti-RBD antibodies. (**E**) Relative abundance of bacterial genera of greater than 1% abundance in sorted bacterial fractions bound by various anti-RBD antibodies. 16S rRNA V3-V4 region of sorted bacteria was sequenced and annotated to corresponding bacteria. Abundance was calculated in relation to the number of total reads. Genera with abundance higher than 1 % were further selected. Frequencies of selected genera were further normalized to 100%. (**F**) List of cloned bacteria isolated based on the binding to anti-RBD antibodies.

To identify the bacteria recognized by neutraliing anti-RBD antibodies, we stained, sorted and sequenced antibody-bound fecal bacteria from 3 healthy donors using 4 different anti-RBD antibodies (Fig. 2D, E). Several genera with an abundance of more than 1% were bound by the respective antibodies, and the identified bacteria differed among various donors (Fig. 2E), highlighting the inter-individual diversity in the bacterial composition. The binding of the anti-RBD IgG antibodies to microbiota was specific, since neither the secondary anti-IgG antibodies used to identify their binding (Fig.2), nor human IgG antibody with different specificity showed similar binding patterns towards microbiota (Fig. S3B). The monoclonal human anti-RBD antibodies in particular showed reactivity towards *Bacteroides*. Some of them also recognised *Clostridia* species, *Streptococci, Escherichia* and *Bifidobacteria* (Fig. 2E). Of the genera bound by IgA of HC RBD-IgA^+^ donors, *Parabacteroides* and *Bilophila* also bound to the human anti-RBD IgG antibodies (Fig. 1I, 2E).

By fluorescence-activated cell sorting we isolated bacteria recognised by the human anti-RBD IgG antibodies from 8 healthy donors, and cultured them using selective bacterial media and anaerobic culture conditions. Individual bacterial colonies were further expanded and their identity determined by 16S rRNA Sanger sequencing (Fig. 2F). Two *Bacilli* species, three *Streptococcus* species, two *Bifidobacterium* species, two *Enterococcus* species, *Veillonella parvula* and *Acidaminococcus intestinalis* were identified as bacteria bound by anti-RBD antibodies (Fig. 2F). Restaining of purified cultures confirmed their recognition by anti-RBD antibodies (Fig. S4A, B). One of the isolated bacterial species was *Streptococcus salivarius*, bacteria living in the oropharynx, with probiotic activity. Indeed, *S. salivarius* K12, an established probiotic strain, is recognized by rabbit anti-RBD antibodies (Fig. S4A). Of note, some bacterial cultures showed only partial staining with anti-RBD antibodies, probably reflecting the heterogeneity of bacteria during growth or community-dependent surface variability. Since the main route of infection with SARS-CoV-2 is via the respiratory tract, we analysed the reactivity of salivary IgA against the oropharyngeal bacteria *S. salivarius K12, B. pseudocatenulatum* and *B. subtilis*. Saliva from HC RBD-IgA^+^ donors contained significant levels of IgA1 and IgA2 binding to *S. salivarius* and *B. pseudocatenulatum* (Fig. S4C). Western blot analysis of bacterial lysates revealed that rabbit anti-RBD antibody and the human anti-RBD IgG antibody HL CV07-200 recognise discrete proteins of *S. salivarius* and *B. pseudocatenulatum* which were further identified by mass-spectrometry (Fig. S5A-E). Subsequent cloning and overexpression in *E. coli* showed binding of anti-RBD antibody to “uncharacterised protein RSSL-01370” of *S. salivarius K12* (Fig. S5B). These data demonstrate that commensal microbiota express distinct protein antigens that are recognized by some, but not all, neutralising anti-RBD antibodies.

Having shown that anti-RBD antibodies can cross-react with bacterial proteins, we tested whether the bacteria expressing these proteins can induce a cross-reactive anti-RBD antibody response. We immunised C57Bl/6 mice intraperitoneally once with heat-killed bacteria and analysed the antibody responses against RBD 14 days later. Mice immunized with heat-killed *S. salivarius*, but not those immunized with heat-killed *B. pseudocatenulatum*, developed anti-RBD IgG antibodies in their sera (Fig. 3A). *Veillonella parvulla* also induced anti-RBD IgG upon immunization (Fig. 3B). Sera from mice immunised with *S. salivariu*s and *V. parvulla* could inhibit the binding of RBD to ACE2, as expressed in 293 T cells (Fig. 3C). Closer to the physiological situation, the natural route of confrontation with bacteria of oropharyngeal microbiota, oral feeding with *S. salivarius K12* and *B. pseudocatenulatum*, induced fecal IgA specific for RBD in C57Bl/6 mice (Fig. 3D). Moreover, fecal supernatants from animals supplemented with bacteria inhibited binding of RBD to ACE2 (Fig. 3E). To gain further insight on the specificity of antibodies induced by oral supplementation with bacteria, we next performed epitope mapping of the IgA induced in the gut against 564 peptides derived from the Spike protein of SARS-CoV-2. We observed that both *B. pseudocatenulatum* and *S. salivarius* induced antibodies bound to the peptide sequence GFNCYFPLQSYGFQPTNGV (Fig. 3F, Fig. S6), that corresponds to the receptor binding motif (RBM) of RBD, in line with ACE2 inhibition data. Also, the peptide recognition pattern of rabbit anti-RBD and HL CV07-200 antibodies overlapped: both antibodies had in their epitopes a similar sequences within the RBM motif (Fig. S6). These data show that oral supplementation with *S. salivarius K12* and *B. pseudocatenulatum* can induce antibodies cross-reactive against the RBM motif of the spike protein of SARS-CoV-2.

**Fig. 3.**
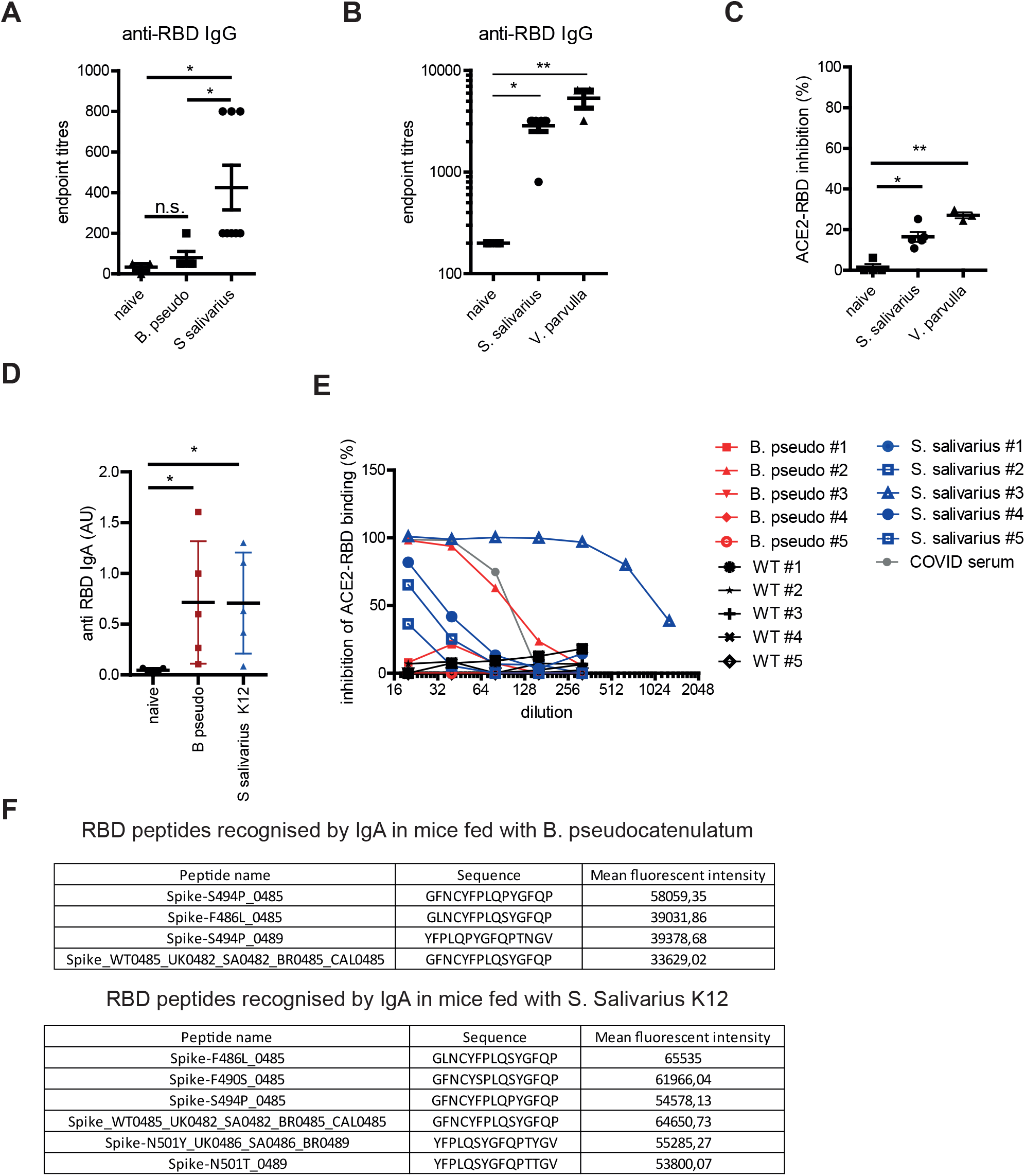
Commensal microbiota species can induce neutralising anti-RBD response. **(A)** anti-RBD IgG titers at day 14 in mice immunized with heat-killed S. salivarius and B. pseudocatenulatum. Mice were immunized as described in materials and methods. **(B)** anti-RBD IgG titers in mice immunized with isolated, heat-killed V. parvulla and S. salivarius K12. (**C**) Inhibition of RBD binding to ACE2 by sera from animals primed with heat-inactivated bacteria 14 days after immunization. (**D**) Induction of anti-RBD IgA response by oral bacterial supplementation. Mice were orally gavaged every second day as described in materials and methods. Anti-RBD IgA was analyzed in fecal supernatants. **(E)** Inhibition of ACE2-RBD binding by fecal supernatants from mice treated as in **D. (F)** RBD peptides recognized by the antibodies elicited upon oral supplementation of mice with S. salivarius K12 and B. pseudocatenulatum for 3 weeks. Kruskal-Wallis test with Dunn’s multiple comparisons was used for (**C**) and (**D**). *, p<0.05, **, p<0.01, ***, p<0.001, ns, not significant. Two-way ANOVA with Bonferroni’s correction was applied for the statistical evaluation of (A) and (B).

In light of the ability of distinct oropharyngeal microbiota species to generate mucosal IgA cross-reactive to SARS-CoV-2, we compared the oral microbiota composition of healthy donors to that of COVID-19 patients, as well as of patients with flu-like symptoms, but negative for SARS-CoV-2 (Fig. 4 and Table S2). A principal component analysis (PCA) indicated that the oral microbiota of hospitalized COVID-19 patients differed considerably from healthy donors, patients with mild COVID-19 and patients with flu-like symptoms (Fig. 4A). First of all, the oral microbiota from severely diseased COVID-19 patients was characterized by an overall decreased bacterial diversity (Fig. 4B). A subsequent LefSE analysis revealed multiple bacterial genera enriched in severe COVID-19 patients (Fig. 4C). Conversely, *Veillonella* and *Streptococcus* genera, but not *Bifidobacteri*a genera, which we had identified as potential inducers of cross-reactive antibodies, were significantly reduced in patients with severe COVID-19 (Fig. 4C, D). Instead, these patients showed an increased abundance of the genera *Enterococcus, Staphylococcus* and *Escherichia/Shigella* in their oropharynx (Fig. 4C, D). This is not due to the treatment of severe COVID-19 patients with antibiotics (Abx), since our cohort includes both Abx naive and Abx-treated patients, and both groups showed the same prevalence of microbiota composition.

**Fig. 4.**
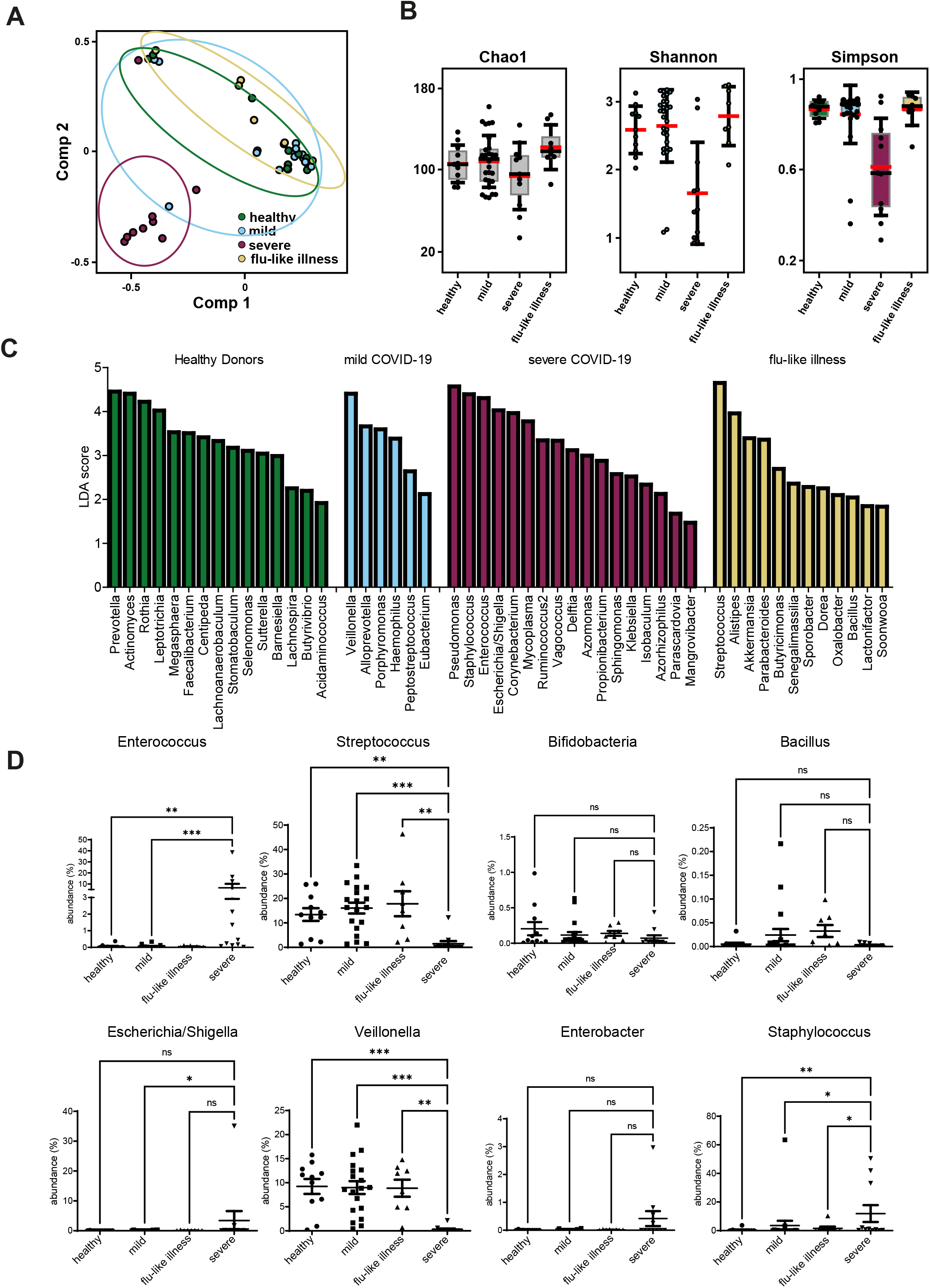
Oral microbiota changes during COVID-19. (**A**) Principal Component analysis of microbiota of swabs collected from healthy individuals, mild, and severe COVID-19 patients and patients with flu-like illness. **(B)** Species richness (Chao1 index) and microbial diversity (Shannon and Simpson index) in oral microbiota collected by swabbing of healthy, mild and severe COVID-19 patients and patients with flu-like illness. **(C)** LDA scores of genera between healthy, mild and severe COVID-19 and flu-like illness. Linear discriminant analysis (LDA) combined with effect size measurements (LEfSe) was performed for 16S rRNA datasets obtained from swabs of healthy, mild and severe COVID-19 and flu-like illness. A p-value of < 0.05 was considered significant in Kruskal–Wallis test and were depicted on the figure. (**D**) Abundance of selected bacterial genera in swabs from healthy, mild and severe COVID-19 and flu-like illness. Kruskal-Wallis test with Dunn’s multiple comparisons was used for (**D**) unpaired t-test was used for (**B**). *, p<0.05, **, p<0.01, ***, p<0.001, ns, not significant.

The differences in the oral microbiota composition also extend to the intestinal microbiota in severely affected COVID-19 patients (Fig. S7A and Table S3). Also intestinal microbiota from severe COVID-19 patients displayed reduced bacterial diversity (Fig. S7B) with dominance of opportunistic pathogenic bacteria, such as *Enterococcus, Staphylococcus* and *Vagococcus* (Fig. S7C, D). *Streptococcus* genera were significantly diminished in severe COVID-19 patients (Fig. S7C, D), while the differences in *Bifidobacteria* genera were not significant. Thus, severe COVID-19 is associated with the outgrowth of opportunistic bacteria (*Enterococci, Staphylococc*i), while other genera, like *Streptococcus* and *Veillonella* are depleted from the mucosal surfaces, both oral and intestinal.

SARS-CoV-2 infection of human mucosal surfaces induces an inflammatory syndrome that may progress towards fatal disease. Multiple factors, both host-intrinsic and host-extrinsic, were uncovered as drivers of disease progression. Host-derived risk factors include presence of autoantibodies against type I IFN, genetic predisposition [30, 31] and preexisting disease conditions, such as diabetes, obesity, and ageing [32]. Furthermore, while pre-existing memory T cells specific for SARS-CoV-2 may be protective, pre-existing low avidity memory T cells recognising SARS-CoV-2 antigens in the elderly may be a potential risk factor during COVID-19 [8, 33]. Here we report that healthy, unexposed individuals can have preexisting secretory IgA antibodies at mucosal surfaces, antibodies which also bind to the RBD of the S protein of SARS-CoV-2, and thus have the potential to neutralise the virus and prevent or ameliorate infection and COVID-19. This pre-existing mucosal immunity fades with age.

Microbiota may contribute to the protection of the host from infection via modulating the ACE2 receptor expression [34], induction of tonic type I IFN responses [35], and via tuning systemic and mucosal TGF-β1 levels, with TGF-β1 being the inductor of antibody class switch recombination to IgA [8, 36]. Here we have identified bacteria of the oropharyngeal microbiota that express protein antigens on their cell surface, which mimic epitopes of the RBD of the SARS-CoV-2 Spike protein, to an extent that they not only are recognised by anti-RBD antibodies of different origin but can themselves also trigger an antibody response capable of neutralising RBD in mice *in vivo*, both by intraperitoneal immunization and by oral feeding. Presence of these bacteria is associated with mucosal IgA antibodies recognizing RBD, and are capable of inhibiting its binding to ACE2, in healthy donors not previously exposed to SARS-CoV-2. It remains a challenge for future research, to determine how the bacteria induce such antibodies. Similar observations have been reported for the HIV-1 virus [27, 37, 38]. In particular, a link between gp-41 and gp-120 reactive antibodies and their cross-reactivity against microbiota has been demonstrated [27, 37]. It is evident that bacteria of the microbiota provide a rich target proteome for the mucosal immune system, and that this can result in the generation of a cross-reactive, pre-existing mucosal immunity against distinct viruses and may explain heterogeneity of human subjects in susceptibility towards viral infection.

Apart from host-intrinsic factors, the initial virus load may affect disease outcome and severity [39, 40], and there is an increasing evidence of microbiota changes during severe COVID-19 [41, 42], suggesting that the microbiota composition may be a risk factor for the development of severe disease as well [41, 42, 43]. The data are conflicting in terms of the genera associated with disease severity, which is probably due to the heterogeneity of patient cohorts and differences in treatment. A common denominator is that acute COVID-19 is associated with the prevalence of opportunistic bacteria and depletion of immunomodulatory bacteria [42]. The present study, showing an increase in *Enterococci, Staphylococci*, and *Vagococci*, and depletion of *Veillonella* and *Streptococc*i species in severe COVID-19 patients, is in line with this notion. But whether these changes are cause or consequence of SARS-CoV-2 infection has remained unclear.

On one hand, our data show that microbiota can be recognised by the antibodies raised against the RBD domain of the SARS-CoV-2 spike protein. Such antibodies presumably also shape the microbiota composition. Further studies analysing the impact of antibody responses induced by virus infection and vaccination induced on the microbiota composition are needed to address this fundamental question. On the other hand, immunocompromised patients and patients using immunosuppressive drugs respond poorly to the vaccination [44]. Data presented here propose that bacteria supplementation, in particular with *S. Salivarius K12*, may enhance the titers of anti-RBD IgA antibodies at the mucosal surfaces, prophylactically or therapeutically, or even in the context of vaccination.

The data presented here propose that bacterial supplementation either prophylactically, therapeutically, or in the context of vaccination, and particularly with *S. Salivarius K12*, may enhance the titers of anti-RBD IgA antibodies at the mucosal surfaces.

### Limitations of the study

Despite of the identification of various bacteria that can induce cross-reactive immune responses against the RBD domain of the SARS-CoV-2 Spike protein, it remains to be determined whether induction of such antibodies in humans may protect from SARS-CoV-2 infection or severe course of COVID-19. Further, the size of the cohorts used in this study, both healthy and COVID-19 patients included in the study does not allow for a detailed correlation analysis of microbiota-induced anti-RBD antibody responses with the outcome of COVID-19.

## Materials and Methods

### Human Donors

The recruitment of study subjects was conducted in accordance with the Ethics Committee of the Charité (EA 1/144/13 with EA 1/075/19, EA 2/066/20) and was in compliance with the Declaration of Helsinki.

### Stool sample preparation

Fresh stool samples of patients and healthy controls were stored on ice or at 4°C before processing within 48 h. The stool was diluted in autoclaved and sterile-filtered PBS (in-house, Steritop® Millipore Express®PLUS 0.22 μm, Cat. No: 2GPT05RE) according to weight in the ratio 100 μg/mL and homogenized by vortex and spatula. The feces solution was then subsequently filtered through 70 μm (Falcon, Cat. No. 352350) and 30 μm filters (CellTrics®, Sysmex, Cat. No. 04-0042-2316) and centrifuged at 4000 x g to pellet the bacterial cells. The supernatant of this centrifugation step was once more centrifuged at 13,000 x g to pellet residual cells. The cell free supernatant was filtered through a 0.22 μm syringe top (Filtropur, Sarstedt Cat. No. 83.1826.001) filter and stored at – 80°C until further use. Pellets of both centrifugation steps were pooled and re-suspended in 10 mL PBS to measure the cell density at 600 nm. For each working stock a cell amount resembling 0.4 OD was stored in 1 mL of a 40 % glycerol in LB medium mixture in Safe Seal 2 mL reaction tubes (Sarstedt, Cat. No. 72.695.500) and transferred to – 80°C.

### Swabs sample preparation

Swabs were prepared for 16 S rRNA sequencing with an adapted protocol of the Quick-DNA™ Fecal/Soil Microbe Miniprep Kit (Zymo Research, Cat. No. D6010). Swabs were obtained from clinics on – 80 °C and kept frozen until further use. The swab stick was either already stored in buffer or Bead Bashing™ buffer was added to cover the swab brush. Up to 750 μL of the buffer solutions where transferred to a BashingBead™ Lysis Tube and rigorously mixed at 13,000 rpm at 37 °C. Following the kits protocol the supernatant was harvested after centrifugation at 13,000 x g for 5 min and once more filtered by an Zymo-Spin™ III-F Filter. The DNA containing solution was then treated with Genomic Lysis Buffer and the containing DNA was put on a DNA binding Zymo-Spin™ IICR Column repeatedly until the entire sample volume was loaded. The bound DNA was washed with DNA Pre-Wash Buffer and g-DNA Wash Buffer. The washed DNA was eluted in 50 μL DNA Elution buffer and once more further purified by filtration through the Zymo-Spin™ III-HRC Filter. 2.5 μL of each of the prepared samples was directly loaded to the amplicon PCR of the Illumina Nextera NGS protocol described in the 16 s rRNA method section.

### 16S rRNA gene sequencing

For 16 S rRNA gene sequencing, we amplified the V3/V4 region directly from the sorted samples (primer sequences: 5’-TCgTCggCAgCgTCAgATgTgTATAAgAgACAgCCTACgggNggCWgCAg-3’ and 5’-gTCTCgTgggCTCggAgATgTgTATAAgAgACAggACTACHVgggTATCTAATCC-3’) with a prolonged initial heating step as described by “16S Metagenomic Sequencing Library Preparation” for the Illumina MiSeq System. After the amplicon the genomic DNA was removed by AmPure XP Beads (Beckman Coulter Life Science Cat. No. A63881) with a 1:1.25 ratio of sample to beads (v/v). Next the amplicons were checked for their size and purity on a 1.5 % agarose gel and if suitable subjected to the index PCR using the Nextera XT Index Kit v2 Set C/D (Illumina, FC-131-2003). After index PCR the samples were cleaned again with AmPure XP Beads (Beckman Coulter Life Science Cat. No. A63881) in a 1: 0.8 ratio of sample to beads (v/v). Samples were then analyzed by capillary gel electrophoresis (Agilent Fragment Analyzer 5200) for correct size and purity with the NGS standard sensitivity fragment analysis kit (Agilent Cat. No. DF-473). Of all suitable samples a pool of 2 nM was generated and loaded to the Illumina MySeq 2500 system.

Raw data were processed and de-multiplexed using MiSeq Reporter Software. Forward and reverse reads were combined using PANDAseq 2.11 with a minimum overlap of 25 bases (PMID:22333067) and classified using “classifier.jar” 2.13 from the Ribosomal Database Project with a confidence cutoff of 50% (PMID: 24288368, PMID: 17586664). The copy number adjusted counts were agglomerated to bacterial genera, rarefied to the smallest size and alpha diversity were estimated using phyloSeq 1.34 (PMID: 23630581). Principle coordinate analysis were performed using Bray–Curtis dissimilarity distance using vegan 2.5-7[45].

The linear discriminant analysis were performed using LEfSe, based on copy number adjusted counts normalized to 1M reads [46]. Raw sequence data were deposited at the NCBI Sequence Read Archive (SRA) under the accession number PRJNA738291.

### Microbiota staining

The frozen microbiota stocks were topped up with 1 mL of autoclaved and sterile-filtered PBS to reduce glycerol toxicity while thawing. Samples were centrifuged at 13,000 xg for 10 min twice, the supernatant removed and the pellets re-suspended in PBS and finally divided into 10 tests. All the stainings of microbiota samples were performed in a DNase containing buffer (PBS/ 0.2 % BSA/25 μg/μL DNase, Sigma Aldrich Cat. No. 10104159001). Staining for human immunoglobulins was performed in 100 μL with 1:50 (v/v) of the detection antibodies: anti-human IgM Brilliant Violet 650 (clone: MHM-88, Biolegend® Cat. No. 314526), anti-human IgG PE/ Dazzle™ 594(clone: HP6017, Biolegend® Cat. No. 409324), anti-human IgA1 Alexa Fluor 647 (clone: B3506B4, Southern Biotech Cat. No. 9130-31), anti-human IgA2 Alexa Fluor 488 (clone: A9604D2, Southern Biotech Cat. No. 9140-30). The samples were incubated for 30 minutes at 4 ° C and directly topped up with 1 mL of a 5 μM Hoechst 33342 solution (Thermo Fischer Scientific Cat. No. 62249) for another 30 min at 4 °C. For the detection of Spike protein-similar structures the samples were first incubated in 50 μL containing 0.5 μg SARS-CoV-2 Spike Neutralizing Antibody (clone: HA14JL2302, Sino Biological Inc. Cat. No: 40592-R001) or Neutralizing Antibody isolated from COVID-19 patients for 15 min at 4 °C then washed with PBS and stained again in 50 μL of the anti-Rabbit Alexa 647 (7,5μg/ml, Jackson ImmunoResearch Cat. No. 111-606-144) or anti-human IgG PE/ Dazzle™ 594 (2μg/ml) which was then topped up with 5 μM Hoechst 33342 solution. After Hoechst 33342 staining samples were washed with PBS and centrifuged at 13,000 x g for 5 min. After removal of supernatant, the samples were re-suspended in PBS/ 0.2 % BSA. The samples were transferred to 5 mL round bottom tubes (Falcon, Cat. No. 352063) for acquisition.

### Microbiota Flow Cytometry

We used a BD Influx® cell sorter for all cytometric investigations of the microbiota samples. The sheath buffer (PBS) for the instrument was autoclaved and sterile filtered (Steritop® Millipore Express®PLUS 0.22 μm, Cat. No: 2GPT05RE) before each fluidics start up. The quality of each acquisition was assured by the alignment of lasers, laser delays and laser intensities by Sphero™ Rainbow Particles (BD Biosciences Cat. No. 559123). For sorting, the drop delay was determined prior with Accudrop Beads (BD Biosciences Cat. No. 345249). Samples were acquired with an event rate below 15,000 events and sorted with an event rate below 10,000 events. We always recorded 300,000 Hoechst 33342 positive events. We sorted up to 100,000 events for sequencing directly into Protein Low Bind tubes (Eppendorf Cat. No 022431102), spun down the sample at 17,000 x g and replaced residual sorting buffer by DEPC treated water (Invitrogen Cat. No. 46-2224). The samples were stored in approx. 10 μL at -20°C until further processing. For subsequent cultivation of bacteria, we sorted directly into PYG medium and transferred the cells directly into a COY anaerobic chamber.

### Bacteria culture

PYG medium and plates were prepared as described by the DSMZ (German Collection of Microorganisms and Cell Cultures). 300,000 events were sorted into 1 ml of PYG medium and directly transferred to a COY anaerobic chamber. Sorted bacteria were plated on PYG, BHI (Brain heart infusion broth, Sigma, Cat. No. 53286-100G) and Fastidious agar plates (Thermo Scientific, Cat. No. 12957138) and bacteria were grown for 24 hours. Colonies were picked and PYG medium, BHI broth and Schaedler broth (Roth, Cat. No. 5772.1) were inoculated with colonies from the respective plates. The next day, DNA was isolated and the remaining bacteria were frozen in 40% glycerol LB medium in liquid nitrogen or – 80 °C.

### Sequencing from bacterial colonies

For the identification of the bacterial species bound to the neutralizing anti-RBD antibodies, the DNA from 200 μl of the grown bacteria was isolated with ethanol precipitation. The isolated DNA was subsequently amplified by the 16S rDNA specific primers LPW57 and LPW58 [47]. In brief, bacterial DNA was amplified with Taq-polymerase (0.005 u/μl, Rapidozym GmbH, Cat. No. GEN-003-1000), 3.12 mM MgCl2 (Rapidozym GmbH), 1 X GenTherm buffer (Rapidozym GmbH), 0.25 mM dNTP mix (Thermo Scientific, Cat. No. R0192) and LPW57 and LPW 58 (1μM, TIB Molbiol) for 35 amplification cycles in a thermocycler. The DNA product was verified by gel electrophoresis and purified with the NuceloSpin Gel and PCR Clean-up Kit (Macherey-Nagel, Cat. No. 740609.50). The concentration of the purified PCR product was adjusted to 5 ng/μl in 15 μl and send to Sanger sequencing by Eurofins Genomics. Sequence identity was determined with the Nucleotide Basic Local Alignment Search Tool (BLAST) provided by NCBI.

### Enzyme-linked immunosorbent assay

For the detection of antibody titers in sera and fecal supernatants 96-well plates were coated with goat anti-human Ig (H+L chain) antibody (Southern Biotech, Cat. No. 2010-01) or goat anti-human IgA Fab (Southern Biotech, Cat. No. 2050-01) antibody for the detection of IgG, IgM and IgA respectively. After washing with 1x PBST for 30 second, the plates were blocked with 200 μL of 5% PBS/BSA for 1 hour at room temperature. Next, plates were washed 3 times with 200 μL of 1x PBST for 30 second at a time. The sera and fecal supernatants were diluted in PBS and 100 μL were added to the plate. Standards were diluted in PBS and applied to the plate: IgA1 (Genway, Cat. No. E04696), IgM (Sigma, Cat. No. 18260), IgA2 (Genway, Cat. No. 50D1F7), IgG (Janssen Biotech Inc.,) then the plates were incubated over night at 4°C. After that, plates were washed 5 times with 200 μL of 1x PBST and detection antibodies were applied: anti-human IgG-AP (ICN/Cappel, Cat No. 59289), anti-human IgM-AP (Sigma, Cat. No.A3437-.25ML), anti-human IgA-AP (Sigma, Cat.No. A2043), anti-human IgA1-AP (SouthernBiotech, clone: B3506B4, Cat. No. 9130-04), anti-human IgA2-AP (SouthernBiotech, clone: A9604D2, Cat. No. 9140-04) and were incubated for 1 hour at 37°C. Subsequently, the plates were washed 5 times with 200 μL of 1x PBST 100 μL of pNPP (Sigma, Cat. No. N2770) was added to each well. Reactions were stopped by addition of 3M NaOH. Optical densities were measured on Spectramax (Molecular devices).

To determine the SARS-Cov-2 specific antibody titers, 96-well plates were coated overnight with either 1 μg/ml recombinant SARS-CoV-2 (2019-nCoV) Spike Protein (RBD, His Tag, Sino biological, Cat. No. 40592-V08B-100) or recombinant SARS-CoV-2 Nucleocapsid His Protein, CF (RnD Systems; Cat. No. 10474-CV) protein or SARS-CoV-2 Spike RBM (receptor binding motif), 480-496 aa (Eurogentec; Cat. No: As-656-19). Plates were washed, blocked and the administration of sera and fecal supernatants were done as previously described **[8]**. To detect RBD-specific IgA, a biotinylated anti-human IgA antibody (Southern Biotech, Cat. No. 2050-08) was applied, followed by an incubation for 1 h at 37°C. After washing 6 times with PBST, avidin-HRP (Invitrogen, Cat. No. 88-7324-88) was added and after 1 hour incubation at RT and 5 times washing with PBST, Tetramethylbenzidine (TMB) Substrate (Invitrogen, Cat. No. 88-7324-88) was added. The reaction was stopped by addition of 2N H2SO4. Optical densities were measured on Spectramax (Molecular devices).

### Epitope mapping for anti-RBD antibodies

Epitope mapping was performed using peptide microarray multiwell replitope SARS-CoV-2 Spike glycoprotein (SPIKE) wild type + mutations (JPT Peptide Technologies GmbH; RT-MW-WCPV-S-V02). Microarray was incubated with monoclonal anti-RBD antibodies (final concentration 1 mcg/ml) or mouse fecal supernatants (1:1 dilution) at 30 C for 1 hours with constant rotation. Slides were washed three times with TBS buffer with 0,05 % Tween-20 and further incubated with anti-rabbit Alexa 647 (Jackson ImmunoResearch Cat. No. 111-606-144), anti-human IgG-Alexa647 (Southern Biotech; Cat. No.: 2040-31), goat anti-Mouse IgA Antibody DyLight® 650 (Bethyl Laboratories; Cat.No.: A90-103D5) at 30 C for 1 hour. Samples were washed with TBS-T and deionized water, dried by centrifugation. Peptide microarray was analysed using microarry scanner Innoscan 710 (Innopsys). Fluorescence intensities were quantified using ImagePix.

### Flow cytometric assay for analysis of ACE2-RBD interaction

HEK293T cells were transfected with a plasmid expressing human ACE2 protein. Next day, the proportion of transfected cells was determined by staining with biotinylated RBD (Sino biologicals, Cat: 40592-V08H-B) for 30 min. The cells were washed s once with PBS/ 0.2 % BSA and subsequently stained with streptavidin-FITC (Thermo Fischer Scientific: Cat. No. 11-4317-87). Further transfected cells were collected and incubated with biological samples for 30 min, washed twice with PBS/BSA and incubated with biotinylated RBD (Sino biologicals, Cat: 40592-V08H-B) for 30 min, washed once with PBS/ 0.2 % BSA and subsequently stained with streptavidin-FITC (Thermo Fischer Scientific: Cat. No. 11-4317-87). Cells were washed with PBS/ 0.2 % BSA measured directly. Dead cell exclusion was done by DAPI. Samples were acquired on a FACSCanto (BD Biosciences) and analyzed using FlowJo v10 (Tree Star Inc.) analysis software.

### Mice immunizations

Grown bacteria were collected, washed three times with PBS and heat-inactivated at 65 C for 1 hr. Heat inactivated bacteria were resuspended with final OD^600^ equals 1.0. C57Bl/6 mice were injected with 200 μl of heat-killed bacteria i.p. From oral gavage, live bacteria stocks were grown, washed with PBS several times, OD^600^ was adjusted to 1, 200 μl of live bacteria was gavaged every second day. All animal procedures were performed in accordance with Russian regulations of animal protection.

### Protein gel electrophoresis and Western blotting

48 h bacterial cultures were pelleted and were resuspended in RIPA buffer containing protease inhibitors cocktail (Roche, Cat. No. 11 836 145 001). Samples were sonicated at 50% voltage for 5 cycles of 10 sec pulses followed by 30 sec rest on ice. After sonication glass beads (MP Biomedicals, Cat. No. 6911100) were added as 1/3 of total volume to the bacterial extract. Samples were vortexed for 30 sec followed by chilling on ice for 30 sec (for a total of 5 cycles). Lysates were spun down for 10 min at 20,000 xg and supernatant was collected. For western blot analysis samples were run on 12% SDS-PAGE under reducing conditions and transferred to PVDF membrane (Bio-Rad, Cat. No. 1620177). SARS-CoV-2 (2019-nCoV) Spike RBD-His Recombinant Protein (Sino Biological, Cat. No. 40592-V08B-100) was used as a positive control. Membrane was blocked by incubation in 5% non-fat milk (Roth, Cat. No. 68514-61-4) in TBST buffer for 1 h at room temperature with constant shaking. Subsequently membrane was hybridized with rabbit neutralizing anti-RBD antibody (Sino Biological, Cat. No. 40592-R001) or human derived RBD neutralising antibodies in blocking solution for 1h at room temperature with constant shaking. Membrane was then washed in TBST and incubated with anti-rabbit IgG-HRP (Cell signaling, Cat. No. 7074S) or with anti-human IgG-HRP (Southern Biotech, Cat. No. 2040-05) for 1 h at room temperature with constant shaking. SuperSignal West Fempto Maximum Sensitivity (Thermo Fisher scientific, Cat No. 34095) substrate kit was used. The signal was acquired using Chemi Doc imaging system (Bio-rad).

### Mass-spectroscopy analysis of proteins

The protein bands after 1D-PAGE were excised and washed twice with 100 mL of 0.1 M NH4HCO3 (pH 7.5) and 50% acetonitrile mixture at 50 °C until the piece of gel becomes transparent. Protein cysteine bonds were reduced with 10mM DTT in 50 mM NH4HCO3 for 30 min at 56 °C and alkylated with 15 mM iodoacetamide in the dark at RT for 30 min. The step with adding DTT was repeated. Then gel pieces were dehydrated with 100 mcl of acetonitrile, air-dried and treated by 10 mkl of 12 mg/mL solution of trypsin (Trypsin Gold, Mass Spectrometry Grade, Promega) in 50 mM ammonium bicarbonate for 15 h at 37°C. Peptides were extracted with 20 mcl of 0.5% trifluoroacetic acid water solution for 30 min with sonication, dried in a SpeedVac (Labconco) and resuspended in 3% ACN, 0.1% TFA. Aliquots (2 mcl) from the sample were mixed on a steel target with 0.3 mcl of 2,5-dihydroxybenzoic acid (SigmaAldrich) solution (30 mg in 400mkl of 30% acetonitrile/0.5% trifluoroacetic acid), and the droplet was left to dry at room temperature. Mass spectra were recorded on the Ultraflex II MALDI-ToF-ToF mass spectrometer (Bruker Daltonik, Germany) equipped with an Nd laser. The [MH]+ molecular ions were measured in reflector mode, the accuracy of the mass peak measurement was 0.007%. Fragment ion spectra were generated by laser-induced dissociation, slightly accelerated by low-energy collision-induced dissociation, using helium as a collision gas. The accuracy of the fragment ions mass peak measurement was 1Da. Correspondence of the found MS/MS fragments to the proteins was performed with the help of Biotools software (Bruker Daltonik, Germany) and a Mascot MS/MS ion search.

### Protein expression

Uncharacterised protein RSSL-01370 was amplified from the genomic DNA of *Streptococcus salivarius K12* using the following primers: 5’-CTCCATATGAATTTACCAAGTCACCATACAAGGG -’3 and 5’-GTGGTCGACATTCACTTTTTCAGTTGCTACACC -’3 and subsequently cloned into pET-21b containing *NdeI* and *XhoI* restriction sites. Next, overnight culture of the selected clone was inoculated into 2xTY growth medium containing 100 μg/ml of ampicillin and grown at 30 °C with constant shaking until OD600 reached 0,8. Protein expression was induced by 0,6 mM of IPTG for the next 4 hours at 30 °C. Bacterial lysate was prepared and analysed as described earlier.

### Purification of IgA from fecal material

Human IgA was purified from fecal supernatants with Peptide M/ Agarose (InvivoGen, Cat. No. gel-pdm-2) as described by the manufacturer. Peptide M/ Agarose was used to prepare a column which was equilibrated with 20 mM sodium phosphate buffer (pH 7). Subsequently the 0.2 μM filtered fecal supernatant was applied on the column at least three times. The fecal IgA was eluted from the column after a washing step with 20 mM sodium phosphate, with 0.1 M Glycine-HCl. The elution was neutralized with 1 M Tris/HCl and was concentrated via dialysis. Finally, the IgA concentration was determined with a NanoDrop 2000C (Thermo Scientific) or ELISA.

## Acknowledgements

We are grateful to Timo Rückert, Lennard Ostendorf, & Marie Burns for the help in blood collection, to the members of the German Rheumatism Research Center Flow Cytometry Core Facility (T. Kaiser, J. Kirsch, & R. Maier) for help with FACS analysis and cell sorting, and to James Cameron for his careful editing. We thank Sergei Nedospasov (MSU, Moscow, Russia) and members of DRFZ B cell club for useful discussions.

## Author contributions

A.K., designed the study. J.N., L. B., M. B. A.K., did most of the experiments and analysed data. J.K., M. R., H.P. generated human monoclonal neutralising anti-RBD antibodies. S.Y. performed IgA purification and analysis of antibody concentrations by ELISA. P.D., G.H. M.-F.M., performed 16S bacteria sequencing. D.M., V.G., I.S., performed mass-spectroscopy of isolated proteins. C.T., S. A., S. T., K. S., P.E., M.W., collected samples, analysed clinical data from COVID-19 patients. M.R., G. S., performed neutralisation assay. H.-D.C., A.R., A.D., M.W., H.P., M.-F.M., A.K. designed the study and contributed to the writing of the manuscript.

