## Supplementary material for "Induction of cross-reactive antibody responses against the RBD domain of the spike protein of SARS-CoV-2 by commensal microbiota": Suppl tables S1-S3; Figure S1-S7

Table S1. Subject Characteristics used for IgA measurement in fecal samples

| Variables | Severe COVID-19 cases | Healthy controls (Fig. 1A) | Healthy controls (Fig. 1C) |
|---|---|---|---|
| Number | 19 | 12 | 19 |
| Male (%) | 14 (74) | 6 (50) | 10 (52,6) |
| Median age, *y* (IQR) | 68 (17) | 60 (8,5) | 29 (13) |
| Comorbidities, n (％) | 6 (40) | 0 (0) | 0 (0) |
| Antibiotic therapy before sampling, n (％) | 7 (37) | 0(0) | 0(0) |
| Tazobactam | 6 (32) | | |
| Piperacillin | 6 (32) | | |
| Clarithromycin | 1 (5) | | |
| Meropenem | 3 (16) | | |
| Vancomycin | 1 (5) | | |
| Cirpofloxacin | 5 (26) | | |
| Ceftazdim | 1 (5) | | |
| Cotrimoxazol | 1 (5) | | |
| Cefotaxim | 1 (5) | | |
| Ampicillin | 1 (5) | | |
| Sulbactam | 1 (5) | | |
| Ertapenem | 1 (5) | | |
| Flucloxacillin | 1 (5) | | |
| Antibiotic therapy during sample collection, n(%) | 2 (11) | | |
| Day after symptoims development, days (IQR) | 17 (24) | | |

NOTE. Values are expressed in number (percentage) and median (interquartile range).

Table S2. Subject Characteristics used for the 16S sequencing of swabs microbiota

| Variables | Severe COVID-19 cases | flu-like controls | ambulant COVID-19 cases | Healthy controls |
|---|---|---|---|---|
| Number | 9 | 8 | 18 | 13 |
| Male (%) | 6 (67) | 4 (50) | 13 (72) | 8 (61) |
| Median age, *y* (IQR) | 58 (18) | 30 (5,5) | 33,5 (14) | 62 (27,5) |
| Comorbidities, n (%) | | 1 (12,5) | 2 (11) | 0 (0) |
| Antibiotic therapy before sampling, n (%) | 4 (44) | 0 (0) | 0 (0) | 0 (0) |
| Tazobactam | 2 (22) | 0 (0) | 0 (0) | 0 (0) |
| Piperacillin | 2 (22) | 0 (0) | 0 (0) | 0 (0) |
| Clarithromycin | 1 (11) | 0 (0) | 0 (0) | 0 (0) |
| Meropenem | 2 (22) | 0 (0) | 0 (0) | 0 (0) |
| Vancomycin | 2 (22) | 0 (0) | 0 (0) | 0 (0) |
| Cirpofloxacin | 3 (33) | 0 (0) | 0 (0) | 0 (0) |
| Ceftazidim/Avibactam | 1 (11) | 0 (0) | 0 (0) | 0 (0) |
| Antibiotic therapy during sample collection, n(%) | 4 (44) | 0 (0) | 0 (0) | 0 (0) |
| Day after symptoims development, days (IQR) | 22 (16,5) | 10 (24) | 10,5 (13,5) | 0 (0) |

NOTE. Values are expressed in number (percentage) and median (interquartile range).

Table S3. Subject Characteristics used for the 16S sequencing of fecal microbiota

| Variables | Severe COVID-19 cases | Healthy controls |
|---|---|---|
| Number | 19 | 13 |
| Male (%) | 14 (74) | 7 (54) |
| Median age, *y* (IQR) | 68 (17) | 60 (9,5) |
| Comorbidities, n (%) | 6 (40) | 0 (0) |
| Antibiotic therapy before sampling, n (%) | 7 (37) | 0(0) |
| Tazobactam | 6 (32) | 0(0) |
| Piperacillin | 6 (32) | 0(0) |
| Clarithromycin | 1 (5) | 0(0) |
| Meropenem | 3 (16) | 0(0) |
| Vancomycin | 1 (5) | 0(0) |
| Cirpofloxacin | 5 (26) | 0(0) |
| Ceftazdim | 1 (5) | 0(0) |
| Cotrimoxazol | 1 (5) | 0(0) |
| Cefotaxim | 1 (5) | 0(0) |
| Ampicillin | 1 (5) | 0(0) |
| Sulbactam | 1 (5) | 0(0) |
| Ertapenem | 1 (5) | 0(0) |
| Flucloxacillin | 1 (5) | 0(0) |
| Antibiotic therapy during sample collection, n(%) | 2 (11) | 0(0) |
| Day after symptoims development, days (IQR) | 17 (24) | 0(0) |

NOTE. Values are expressed in number (percentage) and median (interquartile range).

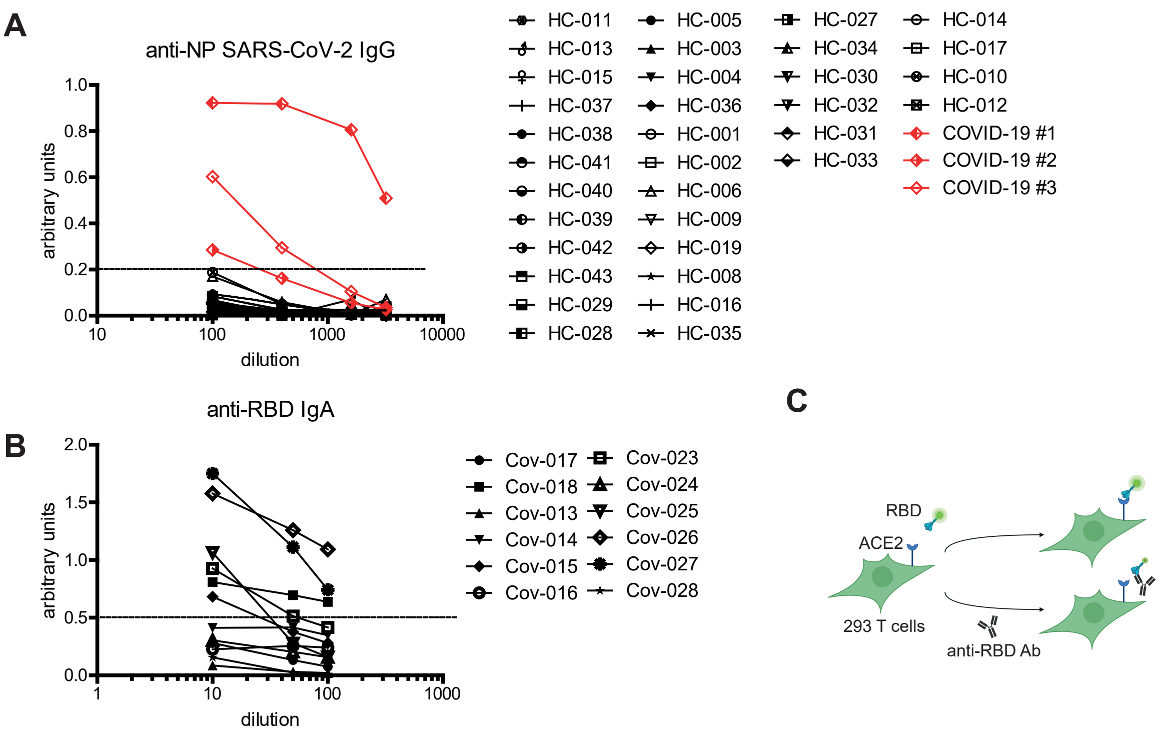


**Figure S1. Characteristics of antibody responses in healthy donors.**
**(A)** Levels of anti-NP SARS-Cov-2 IgG antibodies in sera of healthy donors.
(**B**) Levels of anti-RBD IgA in fecal supernatants of severe COVID-19.
**(C)** Scheme for the analysis of RBD-ACE2 inhibition by flow cytometry.

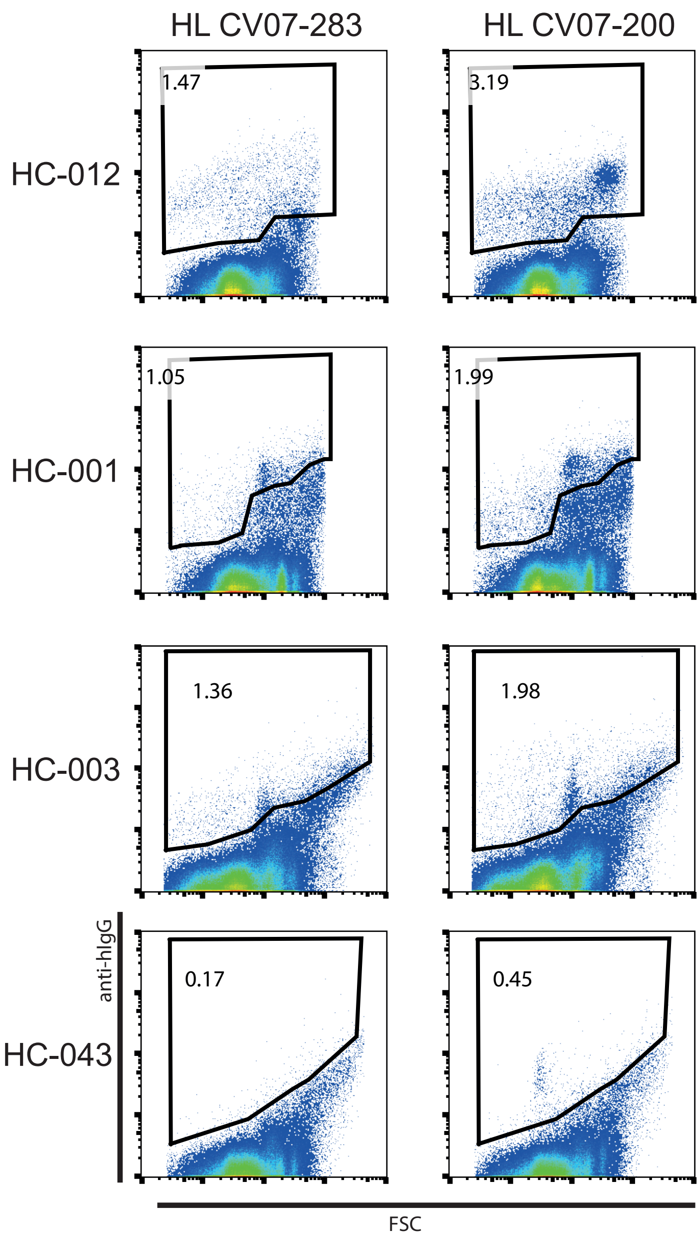


**Figure S2. Binding of clonaly related human anti-RBD antibodies to the microbiota.** Human fecal microbiota was incubated with human anti-RBD antibodies, followed by fluorescently labelled secondary anti-hIgG. Samples were acquired using BD Influx.

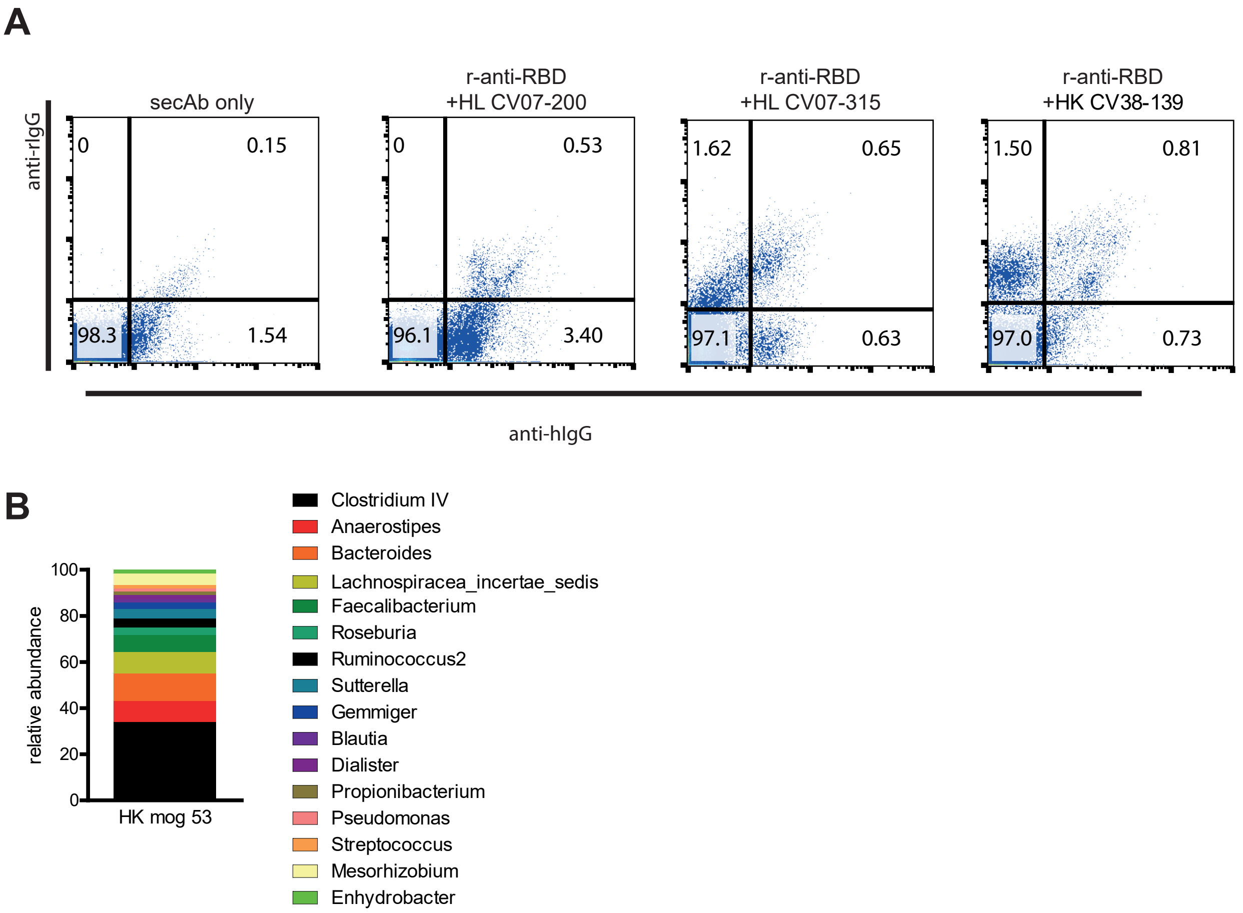


**Figure S3. Recognition of human microbiota by human and rabbit anti-RBD antibodies.** (**A**) Human fecal microbiota was incubated with rabbit anti-RBD and various human anti-RBD antibodies, followed by fluorescently labelled secondary anti-hIgG and anti-rabIgG. (**B**) Relative abundance of bacterial genera of greater than 1% abundance in sorted bacterial fractions bound by HK mog53 antibody. 16S rRNA V3-V4 region of sorted bacteria was sequenced and annotated to corresponding bacteria. Abundance was calculated in relation to the number of total reads. Genera with abundance higher than 1 % were further selected. Frequencies of selected genera were further normalized to 100%. Samples were acquired using BD Influx.

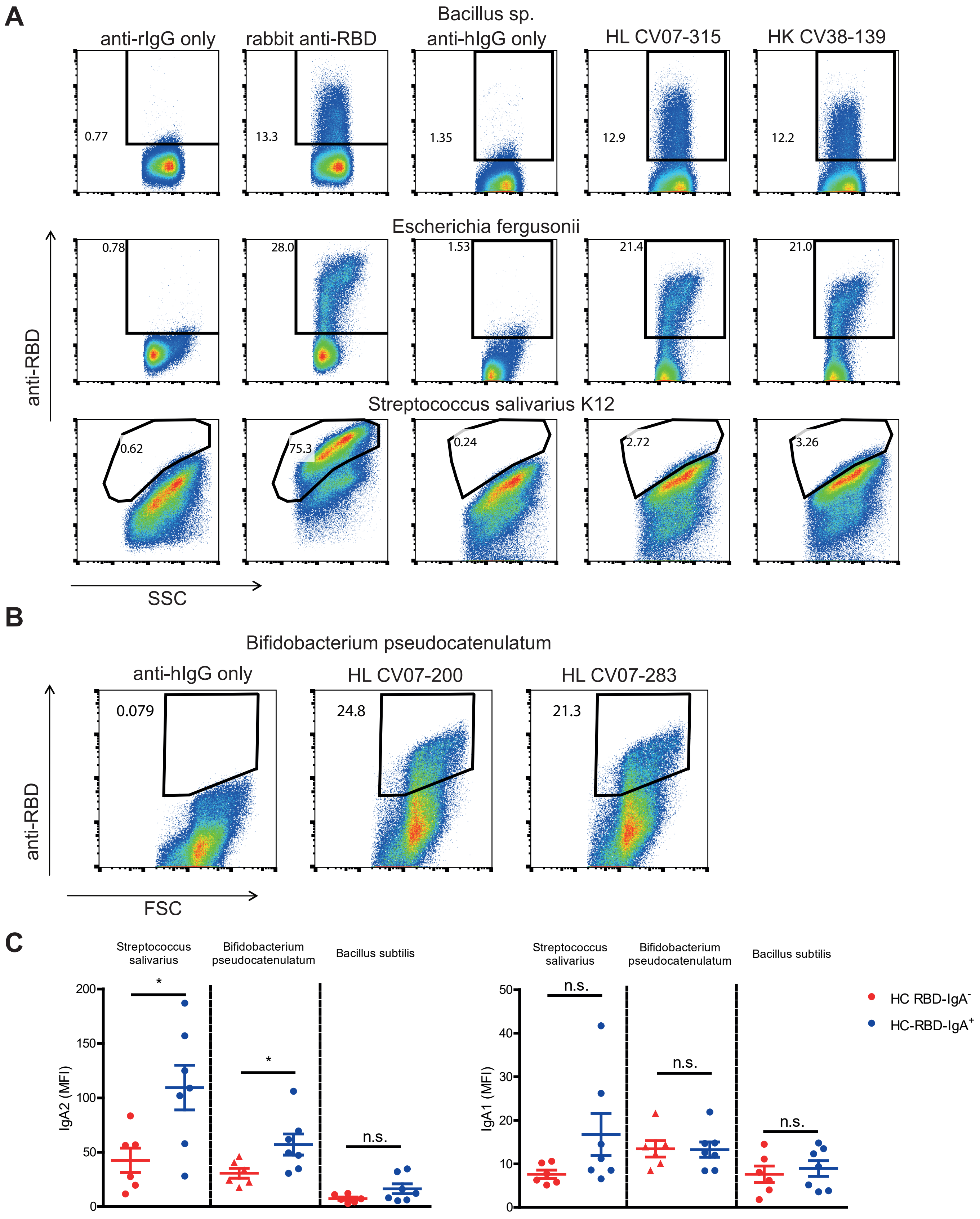


**Figure S4. Recognition of isolated bacterial strains by anti-RBD antibodies.** (**A, B**) Isolated bacterial strains were stained with respective anti-RBD antibodies, followed by fluorescently labelled matching anti-hIgG or anti-rabIgG. Samples were acquired using BD Influx. (**C**) Mean fluorescent intensity of IgA1 and IgA2 coating of various bacterial strains using saliva from HC-RBDIgA$^{+}$ and HC RBD-IgA$^{-}$ donors.

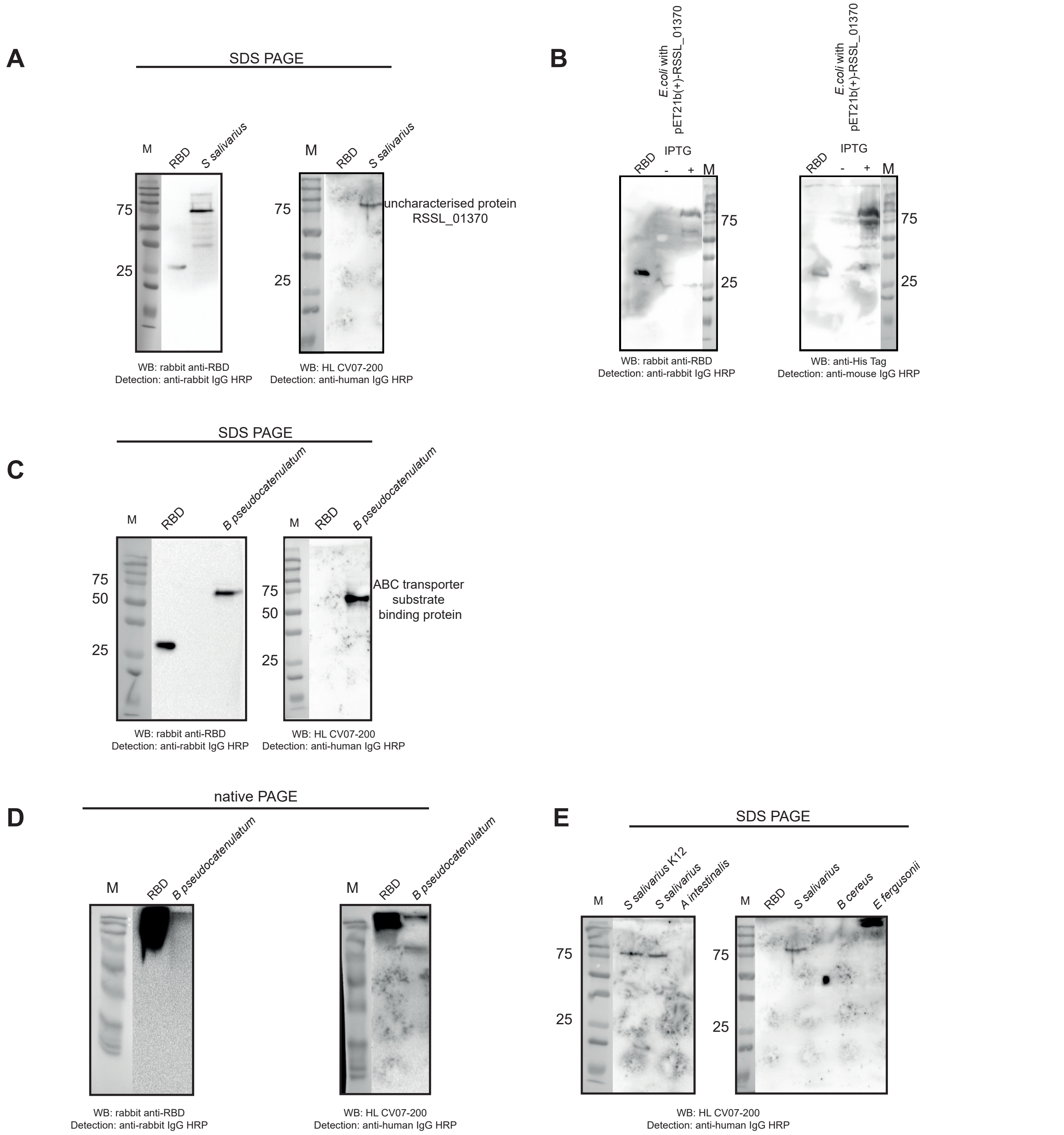


**Figure S5. Anti-RBD antibodies recognise distinct proteins from isolated bacteria.** (**A**) Western blot analysis of rabbit anti-RBD and HL CV07-200 binding to *Streptococcus salivarius* lysate isolated from human microbiota. (**B**) Western blot analysis of rabbit anti-RBD to RSSL_01370 expressed in *E.Coli.* (**C**) Western blot analysis of rabbit anti-RBD and HL CV07-200 binding to *Bifidobacterium pseudocatenulatum* lysate isolated from human microbiota. (**D**) Western blot analysis in native conditions of rabbit anti-RBD and HL CV07-200 binding to *Bifidobacterium pseudocatenulatum* lysate. (**E**) Western blot analysis of HL CV07-200 binding to different bacterial lysates isolated from human microbiota.

**A**

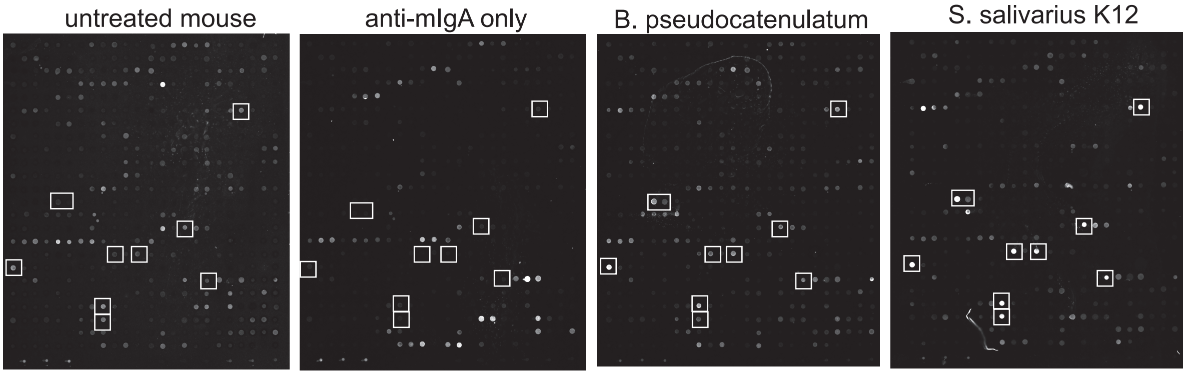


**B**

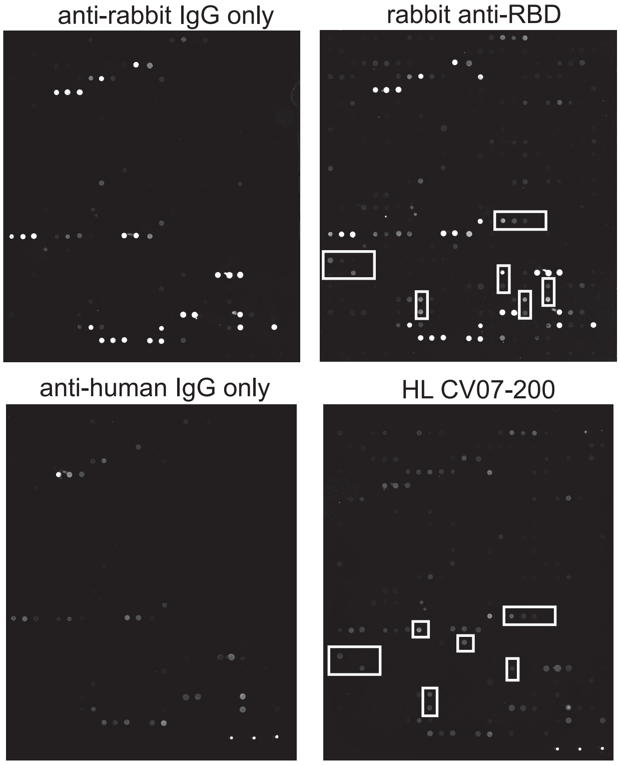


**C**

HL CV07-200

| Peptide Name | Sequence | Mean Fluorescent Intensity |
|---|---|---|
| Spike_WT0325_UK0322_SA0322_BR0325_CAL0325 | SIVRFPNITNLCPFG | 14988,99 |
| Spike_WT0337_UK0334_SA0334_BR0337_CAL0337 | PFGEVFNATRFASVY | 3254,23 |
| Spike_WT0341_UK0338_SA0338_BR0341_CAL0341 | VFNATRFASVYAWNR | 4420,3 |
| Spike_WT0353_UK0350_SA0350_BR0353_CAL0353 | WNRKRISNCVADYSV | 3415,71 |
| Spike-V367F_0353 | WNRKRISNCVADYSF | 5978,49 |
| Spike-L452R_CAL0441 | LDSKVGGNYNYRYRL | 6204,25 |
| Spike-Y453F_0445 | VGGNYNYLFRLFRKS | 5539,81 |
| Spike-Y453F_0453 | FRLFRKSNLKPFERD | 9324,15 |
| Spike-E484K_SA0474_BR0477 | STPCNGVKGFNCYFP | 2911,83 |
| Spike-E484K_SA0478_BR0481 | NGVKGFNCYFPLQSY | 4158,57 |
| Spike_WT0485_UK0482_SA0482_BR0485_CAL0485 | GFNCYFPLQSYGFQP | 8892,57 |
| Spike-F486L_0485 | GLNCYFPLQSYGFQP | 8255,25 |
| Spike-F490S_0485 | GFNCYSPLQSYGFQP | 7954,66 |
| Spike-S494P_0485 | GFNCYFPLQPYGFQP | 5932,7 |
| Spike-N501Y_UK0486_SA0486_BR0489 | YFPLQSYGFQPTYGV | 12468,42 |
| Spike-N501T_0497 | FQPTTGVGYQPYRVV | 3556,91 |
| Spike-N501Y_UK0494_SA0494_BR0497 | FQPTYGVGYQPYRVV | 19061,97 |

**D**

rabbit anti-RBD

| Peptide name | Sequence | Mean fluorescent intensity |
|---|---|---|
| Spike-S494P_0485 | GFNCYFPLQPYGFQP | 8721,37 |
| Spike-F490S_0485 | GFNCYSPLQSYGFQP | 21324,11 |
| Spike-F486L_0485 | GLNCYFPLQSYGFQP | 15343,07 |
| Spike-N501T_0497 | FQPTTGVGYQPYRVV | 47567,9 |
| Spike-N501Y_UK0494_SA0494_BR0497 | FQPTYGVGYQPYRVV | 38947,71 |
| Spike-N501Y_UK0486_SA0486_BR0489 | YFPLQSYGFQPTYGV | 23034,85 |

**Figure S6. Epitope mapping of anti-RBD antibodies.** (**A**) Microarrays were hybridized with fecal supernatants from mice, which were orally gavaged with various bacteria, followed by anti-mIgA-Dylight650 detection. (**B**) Microarrays were hybridized with purified monoclonal antibodies, followed by respective secondary antibodies labelled with Alexa647.
List of peptides bound by HLCV07-200 (**C**) and rabbit anti-RBD (**D**).

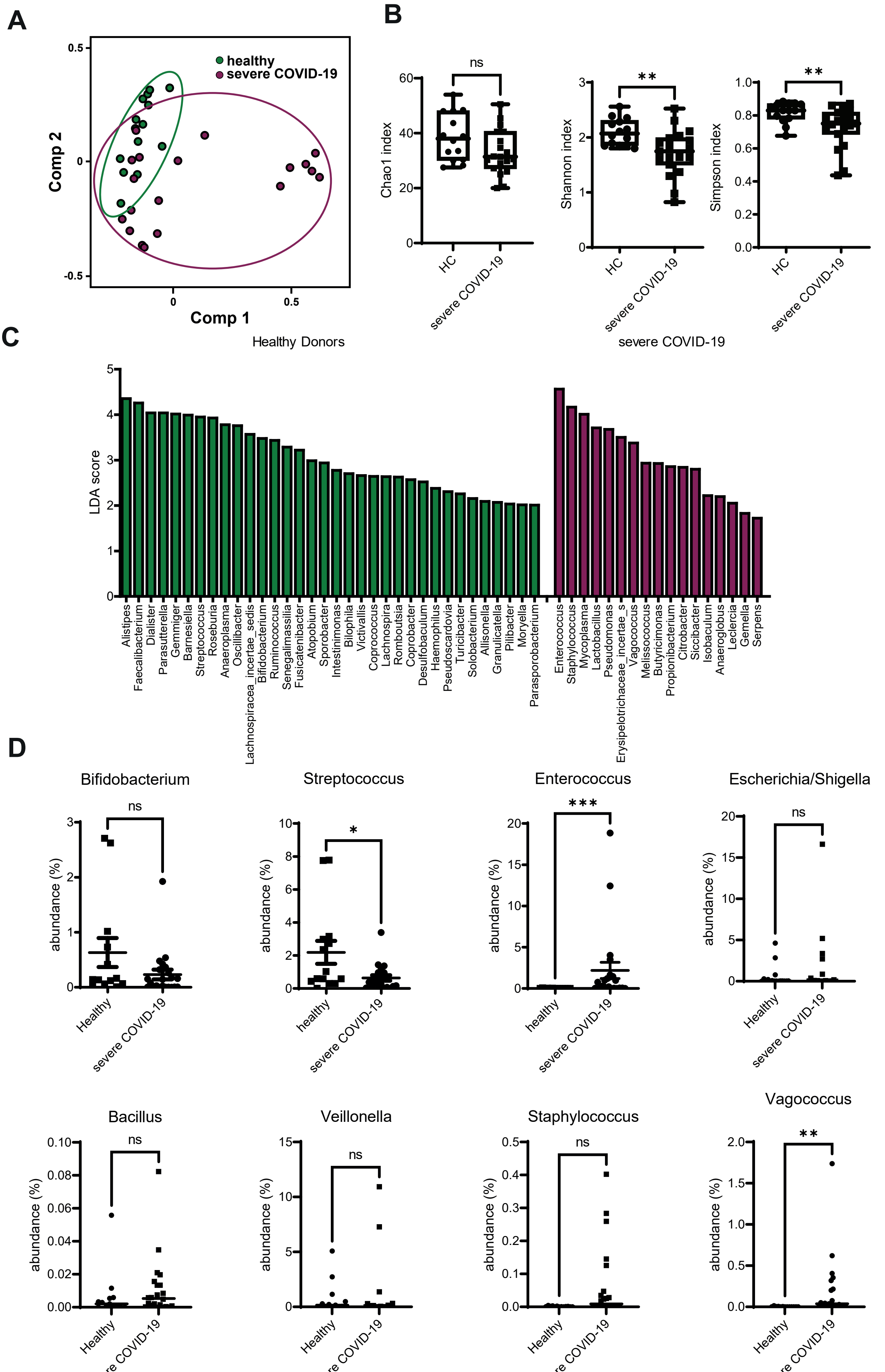


**Figure S7. Fecal microbiota changes during severe COVID-19.** (**A**) Principal Component analysis of microbiota of stools collected from healthy and severe COVID-19 patients. (**B**) Microbial diversity of fecal microbiota of healthy and severe COVID-19 patients. (**C**) LDA scores of genera between fecal microbiota in healthy and severe COVID-19. (**D**) Abundance of selected bacterial genera in feces from healthy and severe COVID-19.
